# Turning a green alga red: engineering astaxanthin biosynthesis by intragenic pseudogene revival in *Chlamydomonas reinhardtii*

**DOI:** 10.1101/535989

**Authors:** Federico Perozeni, Stefano Cazzaniga, Thomas Baier, Francesca Zanoni, Gianni Zoccatelli, Kyle J. Lauersen, Lutz Wobbe, Matteo Ballottari

## Abstract

The green alga *Chlamydomonas reinhardtii* does not synthesize high-value ketocarotenoids like canthaxanthin and astaxanthin, however, a β-carotene ketolase (CrBKT) can be found in its genome. CrBKT is poorly expressed, contains a long C-terminal extension not found in homologues and likely represents a pseudogene in this alga. Here, we used synthetic re-design of this gene to enable its constitutive overexpression from the nuclear genome of *C. reinhardtii.* Overexpression of the optimized CrBKT extended native carotenoid biosynthesis to generate ketocarotenoids in the algal host causing noticeable changes the green algal colour to a reddish-brown. We found that up to 50% of native carotenoids could be converted into astaxanthin and more than 70% into other ketocarotenoids by robust *Cr*BKT overexpression. Modification of the carotenoid metabolism did not impair growth or biomass productivity of *C. reinhardtii*, even at high light intensities. Under different growth conditions, the best performing *Cr*BKT overexpression strain was found to reach ketocarotenoid productivities up to 4.5 mg L^-1^ day^-1^. Astaxanthin productivity in engineered *C. reinhardtii* shown here is competitive with that reported for *Haematococcus lacustris* (formerly *pluvialis*) which is currently the main organism cultivated for industrial astaxanthin production. In addition, the extractability and bio-accessibility of these pigments was much higher in cell wall deficient *C. reinhardtii* than the resting cysts of *H. lacustris*. Engineered *C. reinhardtii* strains could thus be a promising alternative to natural astaxanthin producing algal strains and may open the possibility of other tailor-made pigments from this host.

## Introduction

Carotenoids constitute a widely distributed group of lipid soluble pigments that are synthesized by plants and microorganisms (Kull and Pfander, 1995) and fulfil several important functions in photosynthetic organisms such as light harvesting, light perception, and photoprotection (Mimuro and Katoh, 1991)-(Krieger-Liszkay, 2005). Carotenoids are tetraterpenes, derived from eight isoprene units, (Britton, 1995) containing an extended system of conjugated double bonds that are responsible for their light harvesting and free radical scavenging capacities (Edge et al., 1997). Non-oxygenated carotenoids are named carotenes and this subgroup contains linear (e.g. lycopene) as well as cyclic (e.g. α/β carotene) structures. Oxygenated derivatives of α- and β-carotene are named xanthophylls. Due to their strong colour and anti-oxidant properties, these compounds are widely used in industry as “natural” food colorants, feed additives in aquaculture, and in cosmetics and in pharmaceuticals (Hussein et al., 2006; Li et al., 2011; Yuan et al., 2011). Animals have not been found to synthetize carotenoids naturally, however, they can structurally modify those taken up from their diet (Gerster, 1997).

Among the carotenoids, the secondary ketocarotenoid astaxanthin (3,3’-dihydroxy-β,β-carotene-4,4’-dione) shows superior activity against reactive oxygen species (ROS) and is one of the most powerful natural antioxidants (Naguib, 2000). Astaxanthin synthesis proceeds through oxidation of both rings of β-carotene into canthaxanthin followed by its hydroxylation into astaxanthin (Cunningham and Gantt, 1998; Lotan and Hirschberg, 1995). Alternatively, keto groups can be added to the rings of zeaxanthin, which is derived from the hydroxylation of β-carotene. The enzymes involved in astaxanthin synthesis are 3,3’-β-hydroxylase (*crtz* gene in microalgae) and 4,4’-β-ketolase (BKT, *crtO* gene in microalgae) (Grossman et al., 2004; Lotan and Hirschberg, 1995). Astaxanthin has multiple purported health benefits on biological systems due to its action against ROS (Bennedsen et al., 1999; Jyonouchi et al., 1995). Astaxanthin has potential uses as an antitumor agent (Kim et al., 2016; Palozza et al., 2009; Zhang and Wang, 2015), the prevention of cardiovascular as well as neurological diseases, and diabetes (Gross and Lockwood; Uchiyama et al., 2002; Wu et al., 2015). Moreover, astaxanthin can be used as human dietary supplement and in aquaculture to improve fish color (Hussein et al., 2006; Li et al., 2011; Yuan et al., 2011). Other ketocarotenoids like canthaxanthin, an intermediate of astaxanthin synthesis, has properties similar to astaxanthin, with high potential for use in human health applications (Miki, 1991; Møller et al., 2000). With few exceptions, higher-plants do not synthetize astaxanthin (Cunningham and Gantt, 2011), which is currently produced industrially from unicellular photosynthetic microalgae such as *Haematococcus lacustris* (recently renamed from *H. pluvialis* (Boussiba and Vonshak, 1991; Nakada and Ota, 2016)) or, to a lesser-extent, *Chromochloris zofingiensis* (Chen et al., 2017). *H. lacustris,* is currently the main natural source of astaxanthin as it can accumulate to up to 90% of total carotenoids and 4% of cell dry weight (Bubrick, 1991) under certain environmental conditions. Astaxanthin accumulation in this alga is induced by stress conditions such as nitrogen or phosphorus starvation, high light, salt stress and elevated temperature (Boussiba and Vonshak, 1991) which stimulate the transition from motile zoospores (macrozooids) to immotile spores (aplanospores) (Kobayashi et al., 1997). These changes are accompanied by a degradation of the photosynthetic machinery and cessation of growth (Mascia et al., 2017) as well as the formation of thick and resistant cell walls (cysts) (Boussiba and Vonshak, 1991). The complexities of cellular changes to generate astaxanthin accumulation in *H. lacustris* requires a two-stage cultivation and results in low overall astaxanthin process productivity. Moreover, the recalcitrance of aplanospore cell walls reduce the bio-accessibility of astaxanthin and makes mechanical disruption necessary in order to release astaxanthin for human or animal consumption (Kang and Sim, 2008), a process which increases production process costs.

Given these limitations, genetic engineering approaches have been undertaken to enable astaxanthin production in different biotechnological host organisms in order to generate suitable alternatives to traditional *H. lacustris* production processes. Astaxanthin synthesis has indeed been demonstrated in many different organisms such as fermentative bacteria (Henke et al., 2016; Park et al., 2018) as well as photosynthetic cyanobacteria (Harker and Hirschberg, 1997), and eukaryotic hosts including yeasts (Kildegaard et al., 2017) (Miura et al., 1998) and higher plants (Mann et al., 2000), (Stalberg et al., 2003) (Jayaraj et al., 2008). (Hasunuma et al., 2008), (Zhong et al., 2011) (Huang et al., 2013) (Harada et al., 2014) by the transgenic expression of keto- and hydroxylases. The results obtained were promising but with limited, industrial application due to the high costs of cultivation of these organisms and/or low productivity. Even if high production yield of astaxanthin has been reported upon heterotrophic cultivation of different microorganism, the possibility to produce ketocarotenoids in autotrophic system has the strong advantage in terms of sustainability, consuming CO_2_ and avoiding the costs of reduced carbon sources used in heterotrophic cultivation. In order to develop a sustainable alternative to traditional astaxanthin production, we sought to engineer the common freshwater microalga *Chlamydomonas reinhardtii* to constitutively produce astaxanthin and canthaxanthin. In contrast to previous attempts by others (Leon et al., 2007; Tan et al., 2007; Zheng et al., 2014), our approach is based on the synthetic redesign and revival of an endogenous yet inactive (pseudogene) beta-ketolase sequence present in the nuclear genome of *C. reinhardtii.* Strains resulting from the application of this strategy generated astaxanthin, exhibited reddish-brown phenotypes, and reached productivities comparable to *H. lacustris* cultivation without many of its natural process constraints.

## Results

### Analysis of Chlamydomonas reinhardtii bkt gene

Pathways for the synthesis of carotenoids and xanthophylls in *C. reinhardtii* have already been characterized in previous studies and are depicted in Figure 1 (Lohr et al., 2005). Although, astaxanthin accumulation has never been reported in *C. reinhardtii* in any condition (Lohr et al., 2005), a putative CrBKT enzyme (Uniprot Q4VKB4) can be found in its nuclear genome (Phytozome v5.5). CrBKT has indeed been previously reported to efficiently convert β-carotene and zeaxanthin into astaxanthin when expressed in engineered *E. coli* cells, even in the absence of the CrtZ hydroxylase (Huang et al., 2013; Park et al., 2018; Zhong et al., 2011).

**Figure 1.**
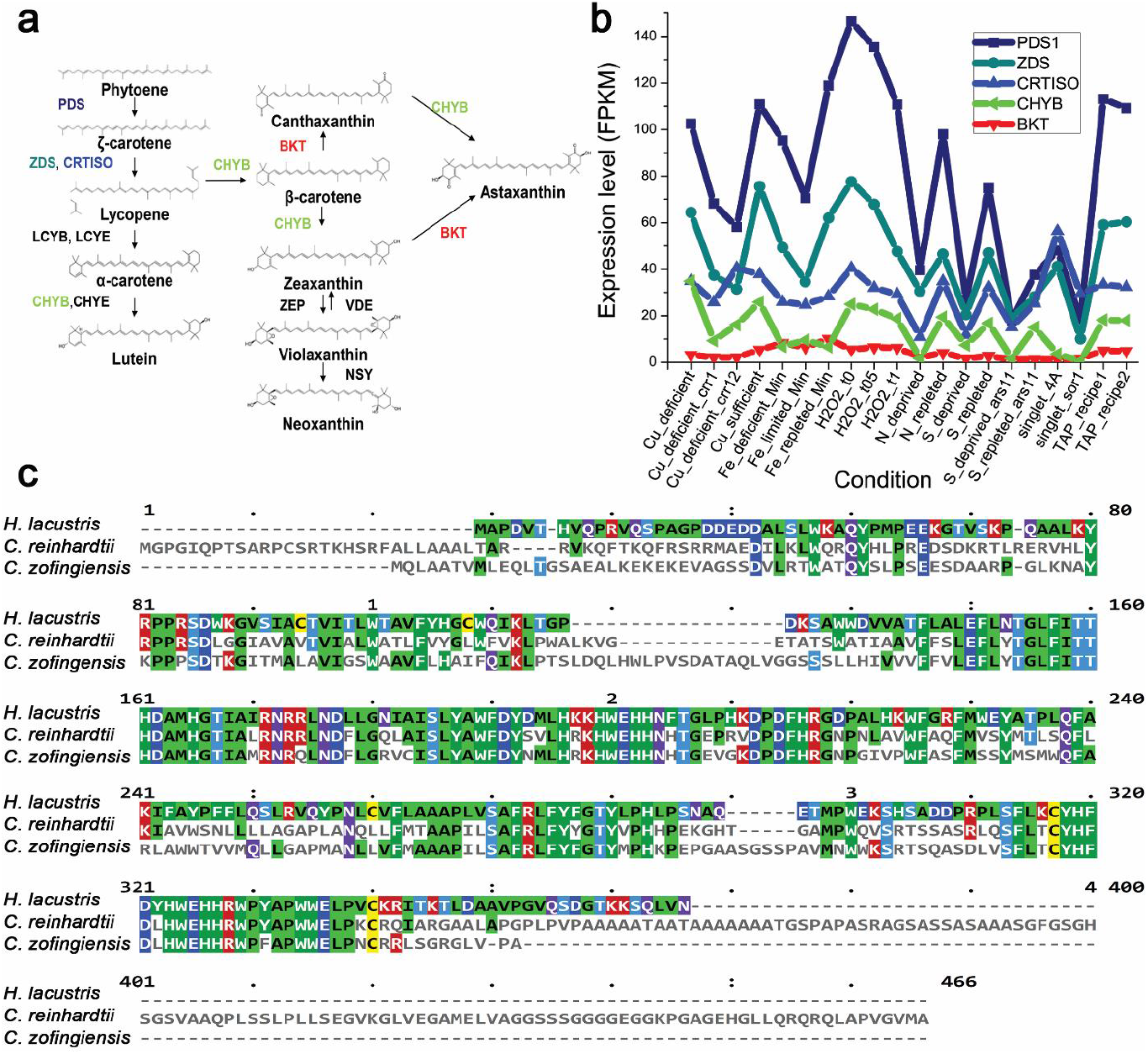
The native BKT of *C. reinhardtii* and its potential role in astaxanthin production. (a) Schematic of the carotenoid pathway towards astaxanthin biosynthesis, according to (Misawa et al., 1995). Only major carotenoids are indicated. Name of enzymes are reported. PDS: phytoene desaturase, ZDS: ζ-carotene desaturase, CRTISO: carotenoid isomerase, LCYB: lycopene β-cyclase, LCYE: lycopene *ε*-cyclase, CHYB:Carotene β-hydoxylase, CHYE: carotene *∈*-hydoxylase, BKT: carotene β-ketolase, ZEP: zeaxanthin epoxidase, VDE: violaxanthin de-epoxidase, NSY: neoxanthin synthase. **(b)** Gene expression level of CrBKT native gene compared to other carotenoid biosynthetic genes. Expression levels in different conditions were retrieved from ChlamyNET database http://viridiplantae.ibvf.csic.es/ChlamyNet/. **(c)**: Protein sequence alignment of BKT from different algae. BKT protein sequences from *C. reinhardtii, H. lacustris* and *C. zofingensis* were aligned by multiple alignment highlighting the long C-terminal amino acid extension of CrBKT. Color code is on the base of consensus.

The native expression level of *CrBKT* gene was thus analysed over a wide range of growth conditions using *C. reinhardtii* RNAseq databases (Romero-Campero et al., 2016). These investigations revealed a very low expression level in any of the tested conditions compared to other genes involved in carotenoid biosynthesis (Figure 1b). An *in-silico* analysis of the *Cr*BKT amino acid sequence revealed an overall high degree of conservation, when compared to other BKT enzymes. A peculiarity of the sequence, however, is the presence of a 116 amino acid C-terminal extension, which is not present in other BKT sequences from other organisms (Lohr et al., 2005) (Figure 1c). A putative chloroplast transit peptide was predicted on the N-terminus of the *Cr*BKT, which was tested for its ability to enable chloroplast import of recombinant mVenus yellow fluorescent protein (hereafter, YFP) in *C. reinhardtii.* While the first 40 N-terminal residues of CrBKT were sufficient for the import of YFP into the chloroplast, a smaller sequence only comprising the first 34 residues from the N-terminus was not (Figure 2a).

**Figure 2.**
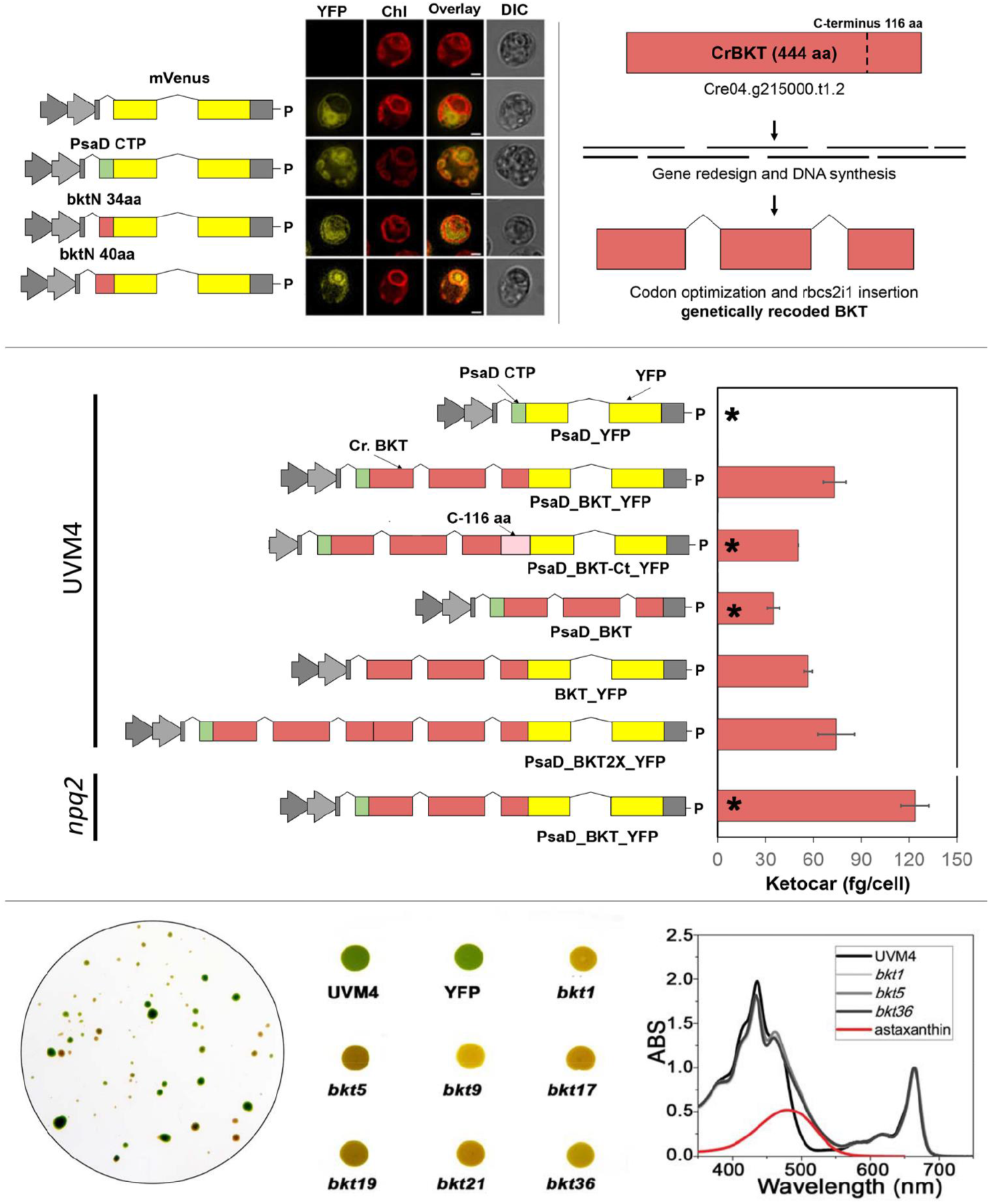
Synthetic BKT redesign, expression vectors for C. *reinhardtii* transformation, and phenotypic change in BKT expression lines. (a) Analysis of endogenous BKT transit peptide was performed by fusion of two different amino acid lengths of the targeting peptide to the YFP reporter. 40 amino acids from the N-terminus of CrBKT was enough to mediate chloroplast import of the reporter. A cytoplasmic and chloroplast targeted control are shown, the latter is mediated by the previously characterized PsaD chloroplast targeting peptide. YFP fluorescence (YFP), chlorophyll autofluorescence (Chl), merger of these two channels and differential interference contrast (DIC) images are shown. Scale bar represents 2 μm. **(b)** The optimized CrBKT sequence was built by gene synthesis after *in silico* design using codon optimization and systematic spreading of the rbcs2 intron 1 sequence to minimize exon lengths as previously described to enable robust transgene expression from the nuclear genome of this alga (Baier et al., 2018b). **(c)** Schematic overview of all expression vectors used in this work and their respective ketocarotenoid accumulation efficiencies. All gene expression cassettes use the Hsp70A/Rbcs2 hybrid promoter containing rbcs2 intron 1 and its 5’ untranslated region as previously described (Lauersen et al., 2015). PsaD_YFP: YFP localized to the chloroplast by the PsaD transit peptide. PsaD_BKT_YFP: BKT fused to YFP targeted by PsaD transit peptide. PsaD_BKT_cterm_YFP_Paro: same as previous vector with the addition of the 116 Cterminal amino acid coding extension of the CrBKT gene. BKT_YFP_Paro: BKT fused to YFP targeted by endogenous transit peptide. PsaD_BKT: CrBKT targeted into chloroplast by PsaD transit peptide without the YFP reporter. PsaD_2xBKT_YFP: Two copies of BKT coding sequence were put in frame in order to generate a fused protein carrying two BKT and YFP. For both *BKT* copies, sequence coding for first 40 aa was removed. All proteins expressed carry a strepII affinity tag (WSHPQFEK*) on the C-terminus. Selection is achieved for all constructs with the AphVIII paromomycin (P) resistance cassette of the pOpt vector backbone. Ketocarotenoid accumulation per cell are expressed as mean ± s.d. (n = 5). The significantly different value from the one obtained with PsaD BKT_YFP construct are marked with an asterisk (*) (P < 0.05). **(d)** Orange/red phenotypes of *C. reinhardtii* cells expressing CrBKT_YFP recovered after transformation and selection on solid medium. **(e)** Image of UVM4 and transformed cells spotted on TAP agar and grown at 100 μmol m^-1^ s^-1^; YFP represents a strain transformed with PsaD_YFP as a control, *bkt* are lines transformed with CrBKT_YFP. (F) Spectra of acetone-extracted pigments from UVM4 and three select *bkt* lines. Spectra are normalized to absorption in Qy region. Spectrum of astaxanthin is shown as reference in red.

### Intragenic expression of β-ketolase in C. reinhardtii

The low native expression rates of CrBKT led us to consider this may be a pseudogene, which has lost activity during evolution and subsequently evolved minimal expression from the algal genome. Given that the gene had been successfully expressed in *E. coli* and shown to functionally convert carotenoids into astaxanthin (Zhong et al., 2011), we decided to investigate whether the gene could be revived by synthetic redesign. Recently, we demonstrated that transgenes could be optimized for expression from the algal nuclear genome by codon optimization and systematic incorporation of the first intron of the *C. reinhardtii* RuBisCO small subunit II gene to mimic host regulatory structures (Baier et al., 2018b). We have recently used this strategy for overexpression of numerous foreign genes (Lauersen et al., 2016; Lauersen et al., 2018; Wichmann et al., 2018), as well as re-coding and overexpression of the endogenous photodecarboxylase (CrFAP) (Yunus et al., 2018) for various metabolic engineering activities in this host. Here, the amino acid sequence of the CrBKT was used to generate an optimized synthetic algal transgene, employing this same optimization strategy (Figure 2b). The optimized gene was first generated by omitting the 116 aa C-terminal extension as this is absent from all other gene homologues in other organisms (Figure 1c) and may influence its activity. The optimized synthetic CrBKT was then cloned into the pOpt2_PsaD_mVenus_Paro vector for expression (Figure 2c) (Wichmann et al., 2018). In order to facilitate chloroplast import, the N-terminus of the BKT coding sequence was fused to the chloroplast transit peptide from photosystem I subunit D (PsaD), which has been already reported to be functional for chloroplast import in different conditions (Lauersen et al., 2015; Rasala et al., 2013). The coding sequence of YFP was left in the vector to generate a fusion at the 3’ end of *BKT.* The construct obtained (PsaD_BKT_YFP, Figure 2c) was then used to transform *C. reinhardtii* UVM4, a parental strain which has been mutated to enable more reliable transgene expression from the nuclear genome (Neupert et al., 2009).

Transformed *C. reinhardtii* recovered on plates exhibited clear changes in colour, from its native green to reddish-brown (Figure 2d). The accumulation of recombinant BKT-YFP protein in transformed cells was then verified by immunoblot developed using an antibody recognizing the fused YFP (Supporting information: Figure S1). *C. reinhardtii* cells expressing *BKT* appeared similar in shape and size under microscopy analysis compared to the background (UVM4), but with a reddish colour (Supporting information: Figure S2). PsaD_BKT_YFP expression lines were screened for the highest accumulation of astaxanthin, first by selection of colonies based on intensity of red colour, then by acetone extraction and spectral analysis (Figure 2e, f). Spectra of pigments from *bkt* lines, extracted with acetone, were found to exhibit a shoulder above 500 nm which was not present in the parental control. This shoulder corresponds to the absorption peak of astaxanthin. Three lines exhibiting the darkest red phenotype and the largest shoulder in spectral analysis were *bkt1, bkt5* and *bkt36.* The content of total ketocarotenoids accumulated in the cells was determined to be up to 73.2 ± 3.7 fg/cell by spectral analysis (see Methods section for further details). In order to possibly improve the expression and activity, further constructs were generated with modifications in the orientation and extensions of the BKT (Figure 2c; UVM4). Many variations of this original expression construct were implemented, however, none resulted in improved astaxanthin production over the PsaD_BKT_YFP vectors (Figure 2c). First, the 116 aa extension removed at the C-terminus of BKT in were reinserted in the vector PsaD_BKT-Ct_YFP, this resulted in an average level of ketocarotenoids ~35% lower compared to the complete version of BKT. The YFP coding sequence was also removed from the C-terminus of BKT (PsaD_BKT), in order to evaluate a possible negative effect due to the presence of YFP at the C-terminus of the protein. The mutants obtained showed a pale red coloration and ketocarotenoids were present although to a much lower level than in the mutants obtained with BKT fused with YFP. A vector coding for a truncated BKT C-terminus fused with YFP and only using the native BKT chloroplast targeting sequence without the PsaD-CTP was also prepared (BKT-YFP) resulting in ~20% lower ketocarotenoid production to the original construct. Finally, two BKT gene copies were fused together in order to enhance the number of catalytic sites (PsaD_BKT2x_YFP) as this strategy had been previously shown beneficial for a sesquiterpene synthase in this host (Lauersen et al., 2016). However, the strains obtained from this construct, showed similar amounts of ketocarotenoids as the single BKT construct (Figure 2c).

The *C. reinhardtii npq2* mutant contains a knockout mutation in the zeaxanthin epoxidase (ZEP, Figure 1a) and is unable to synthetize violaxanthin and neoxanthin, therefore, it accumulates zeaxanthin as a terminal carotenoid species. Zeaxanthin is one of the carotenoid substrates for the BKT enzyme (Zhong et al., 2011). Therefore, to determine if ketocarotenoid yields might be higher in this host, the PsaD_BKT_YFP vector was used to transform the *npq2* mutant (Figure 2c; npq2). After transformation and selection, three lines: *n2bkt1, n2bkt11,* and *n2bkt12,* were selected for strong reddish phenotype and analysed with the same procedure used for *bkt* lines. The ketocarotenoid content per cell of these strains were indeed increased compared to the *bkt* lines obtained in UVM4 background (Figure 2c). Therefore, strains *bkt1/5/36* and *n2bkt1/11/12,* obtained by transformation with construct PsaD_BKT_YFP in the UVM4 and *npq2* backgrounds, respectively, were used for further investigations.

### The presence of ketocarotenoids does not perturb algal growth

The influence of the presence of astaxanthin and ketocarotenoids in C. reinhardtii was evaluated by cultivating selected transformants lines and their parental strains (UVM4, *npq2*) in 20 ml flasks for one week in photoautotrophy (CO_2_) or mixotrophy (acetate) (Harris and Harris, 2008), at 100 or 500 μmol m^-2^ s^-1^. Although *bkt* and *n2bkt* lines exhibited reddish-brown phenotypes, they were nevertheless able to grow even in photoautotrophic conditions (Figure 3a). Both *bkt* and *n2bkt* lines are indeed photosynthetically active and exhibited similar quantum yield of Photosystem II (Fv/Fm) ratios as their parental strains (Figure 3b). Fv/Fm of *npq2* was lower compared to UVM4 which is consistent with previous findings owing to the overaccumulation of zeaxanthin in this strain (Couso et al., 2012). Cell dry weight was measured at stationary phase for all the genotypes (Table 1). At either light intensity tested, growth in mixotrophy was faster compared to the autotrophic cultivation for any strain tested (Figure 3c and e). In both TAP and HS medium, growth of *npq2* strains were slower compared to UVM4, even if the final biomass harvested was similar. Transformant lines exhibited biomass accumulation similar to their respective parental lines. Similar photoautotrophic biomass accumulation in stains containing astaxanthin and those devoid of it, indicated that the presence of astaxanthin does not impair photosynthetic activity.

**Figure 3.**
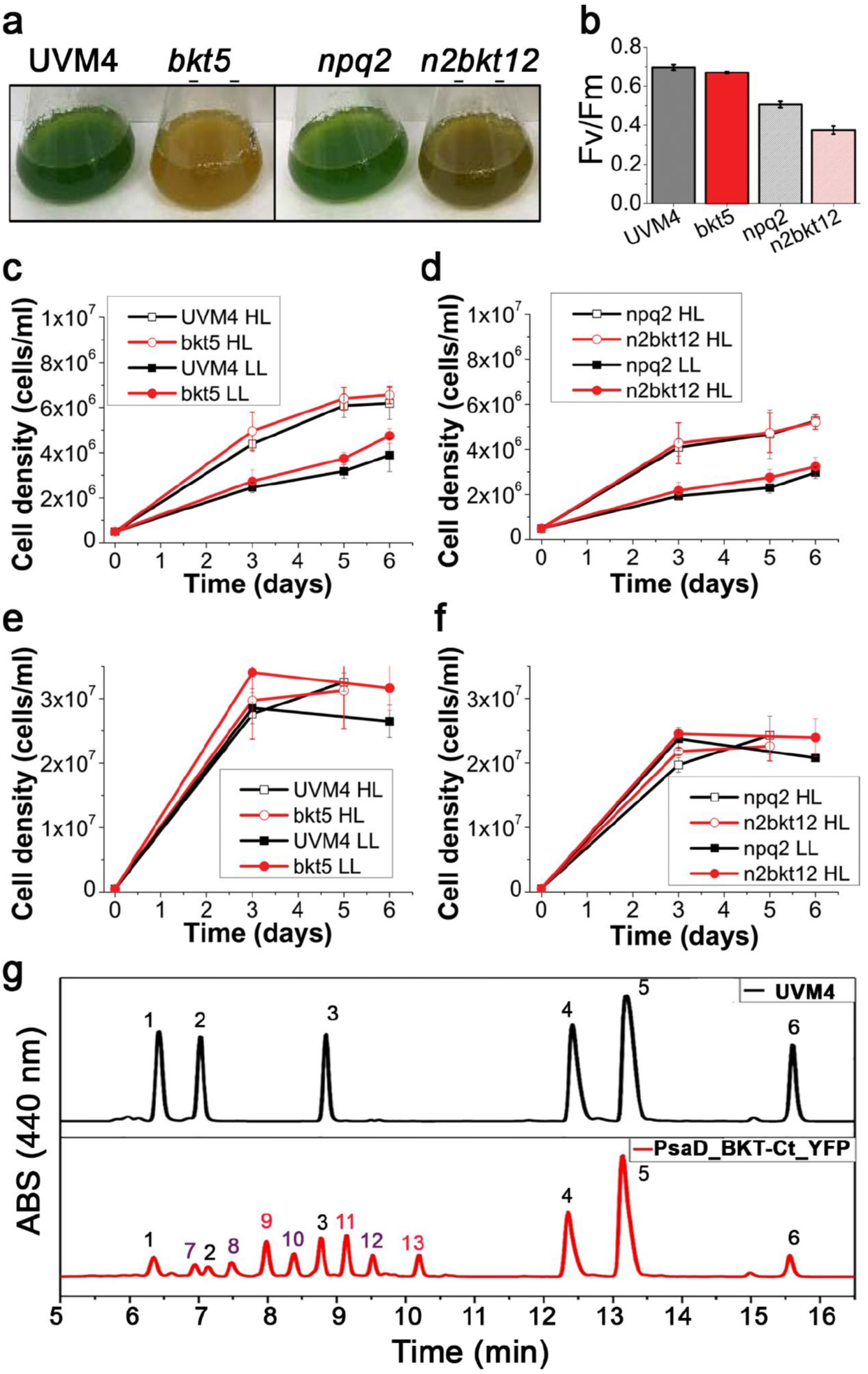
Impact of astaxanthin accumulation in *C. reinhardtii* UVM4 and *npq2* backgrounds on growth. (a) Cultures of UVM4, *bkt5, npq2* and *n2bkt12* grown in autotrophy at 100 umol m^-1^ s^-1^ exhibit striking differences in colour. **(b)** Fv/Fm of UVM4, *bkt5, npq2* and *n2bkt12* grown in autotrophy at 100 umol m^-1^ s^-1^. **(c)** Growth curves of UVM4 and *bkt5* strain in autotrophy (HS medium) or mixotrophy (TAP medium) in low light (LL, 100 umol m^-1^ s^-1^) or high light (HL, 500 umol m^-1^ s^-1^) conditions. **(d)** Growth curves of *npq2* and *n2bkt5* strain in autotrophy (HS medium) or mixotrophy (TAP medium) in low light (LL, 100 umol m^-1^ s^-1^) or high light (HL, 500 umol m^-1^ s^-1^) conditions. **(e)** Representative HPLC of pigments from UVM4 (black) and mutant *bkt5* (red). 1: neoxanthin, 2: violaxanthin; 3: lutein; 4: chlorophyll b; 5: chlorophyll a; 6 β-carotene; 7: unidentified; 8, 10, 12: cantaxanthin; 9, 11, 13: astaxanthin.

**Table 1.** Pigment contents of UVM4, npq2 and *bkt* and *n2bkt* lines. Pigment contents determined in cells grown at 100 or 500 μmol photons m^-1^ s^-1^ in HS and TAP medium for 1 week starting from 5*10^5^ cells/ml. Acronyms indicate: Chlorophyll (chl), total carotenoid (car), astaxanthin (asta), total ketocarotenoid (keto), neoxanthin (neo), violaxanthin (viola), zeaxanthin (zea), β carotene (β car). Chlorophyll per cell are expressed as picograms (pg), astaxanthin and ketocarotenoids as femtograms (fg). Single carotenoids are normalized to 100 chlorophyll molecules. Data are expressed as means ± SD (n = 4). Values marked with the same letters do not differ significantly (p < 0.05).

| growth | genotype | chl pg/cell | chl/car | asta/car | keto/car | asta fg/cell | keto fg/cell | dry weight (g/l) | car/100 chl |  |  |  |  |  |  |
| --- | --- | --- | --- | --- | --- | --- | --- | --- | --- | --- | --- | --- | --- | --- | --- |
| | | | | | | | | | neo | viola | lute | zea | $\beta$ car | asta | keto |
| 100 HS | UVM4 | 0,95 ± 0,09<sup>a</sup> | 2,70 ± 0,21<sup>a</sup> | - | - | - | - | 0,14 ± 0,01<sup>a</sup> | 3,8 ± 0,5<sup>a</sup> | 7,5 ± 1,3<sup>a</sup> | 21,4 ± 3,9<sup>a</sup> | - | 4,3 ± 0,2<sup>a</sup> | - | - |
| | bkt | 0,19 ± 0,04<sup>b</sup> | 1,81 ± 0,14<sup>b,c</sup> | 0,38 ± 0,01<sup>a</sup> | 0,75 ± 0,02<sup>a</sup> | 27,2 ± 2,6<sup>a</sup> | 51,7 ± 5,7<sup>a</sup> | 0,19 ± 0,02<sup>b</sup> | 1,5 ± 0,2<sup>b</sup> | 1,2 ± 0,2<sup>b</sup> | 8,9 ± 1,5<sup>b</sup> | - | 2,1 ± 0,1<sup>b</sup> | 21,3 ± 2,3<sup>a</sup> | 41,6 ± 4,1<sup>a</sup> |
| | npq2 | 0,67 ± 0,20<sup>c</sup> | 2,49 ± 0,67<sup>b</sup> | - | - | - | - | 0,15 ± 0,02<sup>a</sup> | - | - | 18,6 ± 4,5<sup>a</sup> | 16,0 ± 4,3<sup>a</sup> | 5,6 ± 0,5<sup>c</sup> | - | - |
| | n2bkt | 0,25 ± 0,07<sup>b</sup> | 1,42 ± 0,38<sup>c</sup> | 0,41 ± 0,06<sup>a</sup> | 0,76 ± 0,09<sup>a</sup> | 48,3 ± 8,6<sup>b</sup> | 87,2 ± 11,4<sup>b</sup> | 0,14 ± 0,03<sup>a</sup> | - | - | 11,3 ± 2,5<sup>b</sup> | 3,5 ± 0,9<sup>b</sup> | 1,8 ± 0,2<sup>d</sup> | 30,7 ± 10,5<sup>a</sup> | 56,8 ± 18,3<sup>a</sup> |
| 100 TAP | UVM4 | 1,41 ± 0,34<sup>a</sup> | 2,81 ± 0,11<sup>a</sup> | - | - | - | - | 0,56 ± 0,05<sup>a</sup> | 4,9 ± 2,1<sup>a</sup> | 8,7 ± 2,9<sup>a</sup> | 16,8 ± 1,1<sup>a</sup> | - | 5,3 ± 2,1<sup>a</sup> | - | - |
| | bkt | 0,38 ± 0,08<sup>b</sup> | 2,39 ± 0,09<sup>b</sup> | 0,42 ± 0,06<sup>a</sup> | 0,69 ± 0,03<sup>a</sup> | 45,2 ± 6,9<sup>a</sup> | 73,2 ± 3,7<sup>a</sup> | 0,65 ± 0,01<sup>b</sup> | 1,5 ± 0,6<sup>b</sup> | 2,1 ± 0,6<sup>b</sup> | 8,6 ± 0,5<sup>b</sup> | - | 1,0 ± 0,3<sup>b</sup> | 17,4 ± 2,4<sup>a</sup> | 28,8 ± 1,3<sup>a</sup> |
| | npq2 | 1,26 ± 0,49<sup>a</sup> | 2,80 ± 0,03<sup>a</sup> | - | - | - | - | 0,69 ± 0,04<sup>b</sup> | - | - | 12,3 ± 1,3<sup>c</sup> | 19,3 ± 2,0<sup>a</sup> | 4,0 ± 1,2<sup>a</sup> | - | - |
| | n2bkt | 0,72 ± 0,09<sup>c</sup> | 2,45 ± 0,03<sup>b</sup> | 0,3 ± 0,01<sup>b</sup> | 0,62 ± 0,02<sup>b</sup> | 62,1 ± 1,1<sup>b</sup> | 123,7 ± 0,8<sup>b</sup> | 0,65 ± 0,01<sup>b</sup> | - | - | 13,1 ± 1,2<sup>c</sup> | 1,6 ± 0,2<sup>b</sup> | 0,6 ± 0,2<sup>b</sup> | 12,5 ± 0,4<sup>b</sup> | 25,5 ± 0,8<sup>b</sup> |
| 500 HS | UVM4 | 0,87 ± 0,21<sup>a</sup> | 2,54 ± 0,24<sup>a</sup> | - | - | - | - | 0,14 ± 0,01<sup>a</sup> | 3,7 ± 0,9<sup>a</sup> | 7,5 ± 1,8<sup>a</sup> | 21,2 ± 8,3<sup>a</sup> | 2,8 ± 0,4<sup>a</sup> | 4,2 ± 1,1<sup>a</sup> | - | - |
| | bkt | 0,09 ± 0,02<sup>b</sup> | 0,93 ± 0,09<sup>b</sup> | 0,44 ± 0,02<sup>a</sup> | 0,71 ± 0,08<sup>a</sup> | 27,6 ± 2,8<sup>a</sup> | 44,9 ± 10,7<sup>a</sup> | 0,13 ± 0,06<sup>a</sup> | 2,5 ± 0,5<sup>a</sup> | 2,0 ± 0,5<sup>b</sup> | 16,0 ± 5,7<sup>a</sup> | 3,9 ± 0,7<sup>a,b</sup> | 2,3 ± 1,0<sup>b</sup> | 49,1 ± 4,5<sup>a</sup> | 81,1 ± 12,9<sup>a</sup> |
| | npq2 | 0,60 ± 0,24<sup>a</sup> | 2,12 ± 0,73<sup>a</sup> | - | - | - | - | 0,18 ± 0,04<sup>a</sup> | - | - | 20,2 ± 8,0<sup>a</sup> | 22,0 ± 4,2<sup>c</sup> | 4,9 ± 1,6<sup>a,c</sup> | - | - |
| | n2bkt | 0,08 ± 0,03<sup>b</sup> | 0,66 ± 0,22<sup>b</sup> | 0,40 ± 0,07<sup>a</sup> | 0,77 ± 0,04<sup>a</sup> | 56,9 ± 9,4<sup>b</sup> | 106,0 ± 8,4<sup>b</sup> | 0,14 ± 0,03<sup>a</sup> | - | - | 17,2 ± 7,4<sup>a</sup> | 4,7 ± 2,1<sup>b</sup> | 2,8 ± 0,4<sup>b,c</sup> | 44,2 ± 10,1<sup>a</sup> | 84,2 ± 16,5<sup>a</sup> |
| 500 TAP | UVM4 | 1,04 ± 0,17<sup>a</sup> | 2,51 ± 0,34<sup>a</sup> | - | - | - | - | 0,71 ± 0,05<sup>a</sup> | 4,8 ± 1,1<sup>a</sup> | 10,4 ± 2,3<sup>a</sup> | 17,7 ± 3,2<sup>a</sup> | 1,6 ± 0,6<sup>a,b</sup> | 5,4 ± 1,6<sup>a</sup> | - | - |
| | bkt | 0,21 ± 0,03<sup>b</sup> | 1,80 ± 0,24<sup>b</sup> | 0,5 ± 0,04<sup>a</sup> | 0,72 ± 0,02<sup>a</sup> | 39,5 ± 3,3<sup>a</sup> | 56,3 ± 6,9<sup>a</sup> | 0,68 ± 0,03<sup>a,b</sup> | 1,7 ± 0,4<sup>a</sup> | 1,8 ± 0,3<sup>b</sup> | 9,6 ± 1,5<sup>b</sup> | 1,3 ± 0,4<sup>a</sup> | 1,2 ± 0,3<sup>b</sup> | 28,3 ± 5,5<sup>a</sup> | 40,5 ± 5,5<sup>a</sup> |
| | npq2 | 0,98 ± 0,16<sup>a</sup> | 2,39 ± 0,29<sup>a</sup> | - | - | - | - | 0,64 ± 0,07<sup>b,c</sup> | - | - | 14,7 ± 2,4<sup>c</sup> | 22,2 ± 5,4<sup>c</sup> | 5,0 ± 0,2<sup>a</sup> | - | - |
| | n2bkt | 0,47 ± 0,07<sup>c</sup> | 1,67 ± 0,20<sup>b</sup> | 0,41 ± 0,04<sup>b</sup> | 0,7 ± 0,07<sup>a</sup> | 78,7 ± 9,9<sup>b</sup> | 131,7 ± 15,6<sup>b</sup> | 0,62 ± 0,02<sup>c</sup> | - | - | 14,7 ± 2,2<sup>c</sup> | 2,2 ± 0,5<sup>b</sup> | 0,9 ± 0,1<sup>b</sup> | 24,9 ± 5,6<sup>a</sup> | 42,5 ± 8,9<sup>a</sup> |

### Yield of astaxanthin in different growth conditions

Pigments accumulated by *bkt* and *n2bkt* lines were further analysed by high-performance liquid chromatography (HPLC) to verify the accumulation of astaxanthin and other ketocarotenoids. HPLC chromatograms of UVM4 parental strain contains prominent peaks of chlorophylls a and *b* as well as of the carotenoids neoxanthin, violaxanthin, lutein and β carotene (Figure 3g). The transformed lines exhibit additional peaks corresponding to astaxanthin and other ketocarotenoids (mainly canthaxanthin as reported in Figure 3g). Astaxanthin is the major carotenoid in lines expressing PsaD_BKT_YFP, it is present in free, monoester, and diester forms (Figure 3g). In UVM4, ketocarotenoids were never detected while in *bkt* lines, they represent up to 70% of the total carotenoids, with astaxanthin as the major compound (Table 1). Interestingly, in *bkt* lines in all the conditions tested, BKT activity reduced the levels of other carotenoids. Violaxanthin underwent the most pronounced reduction of ~ 80%, β-carotene and neoxanthin were reduced by 65% and lutein 45%. Zeaxanthin was not decreased in *bkt* lines although the amount of this carotenoid was low even in the parental strain. (Table 1). *npq2* lines expressing BKT exhibited strong reductions in zeaxanthin and β-carotene contents, being both substrates from BKT enzyme. In both, *bkt* and *n2bkt* lines, a clear decrease of the chlorophyll amount was evident with the strongest effect in autotrophy and high light conditions (in HS medium at 500 μmol m^-2^ s^-1^, Table 1) This result suggests that upon xanthophyll and β-carotene reduction in favour of astaxanthin accumulation destabilizes the chlorophyll content of photosynthetic complexes, similar to findings previously reported for *H. lacustris* (Mascia et al., 2017). The highest ketocarotenoid content per cell was detected in *n2bkt* lines grown in mixotrophy at 100 or 500 μmol m^-2^ s^-1^, while mixotrophic growth at 500 μmol m^-2^ s^-1^ promoted the highest values in *bkt* lines (Table 1). To further measure productivity of mutants expressing BKT in different conditions, one line with the highest level of astaxanthin for each background, *bkt5* and *n2bkt12,* was selected for astaxanthin productivity analysis in in small scale (80 ml) airlift photobioreactors. These two strains were cultivated at different light regimes (100 and 500 μmol m^-2^ s^-1^), in autotrophy (HS medium) or mixotrophy (TAP medium) and bubbling with air or 3% CO_2_. These experiments were performed in a semi-continuous manner, with harvesting and culture dilution when stationary phase was reached (Figure 4). The experiment was continued until three cycles of dilution were repeated and the productivity of astaxanthin and ketocarotenoids was quantified as mg of pigments per litre per day of culture.

**Figure 4.**
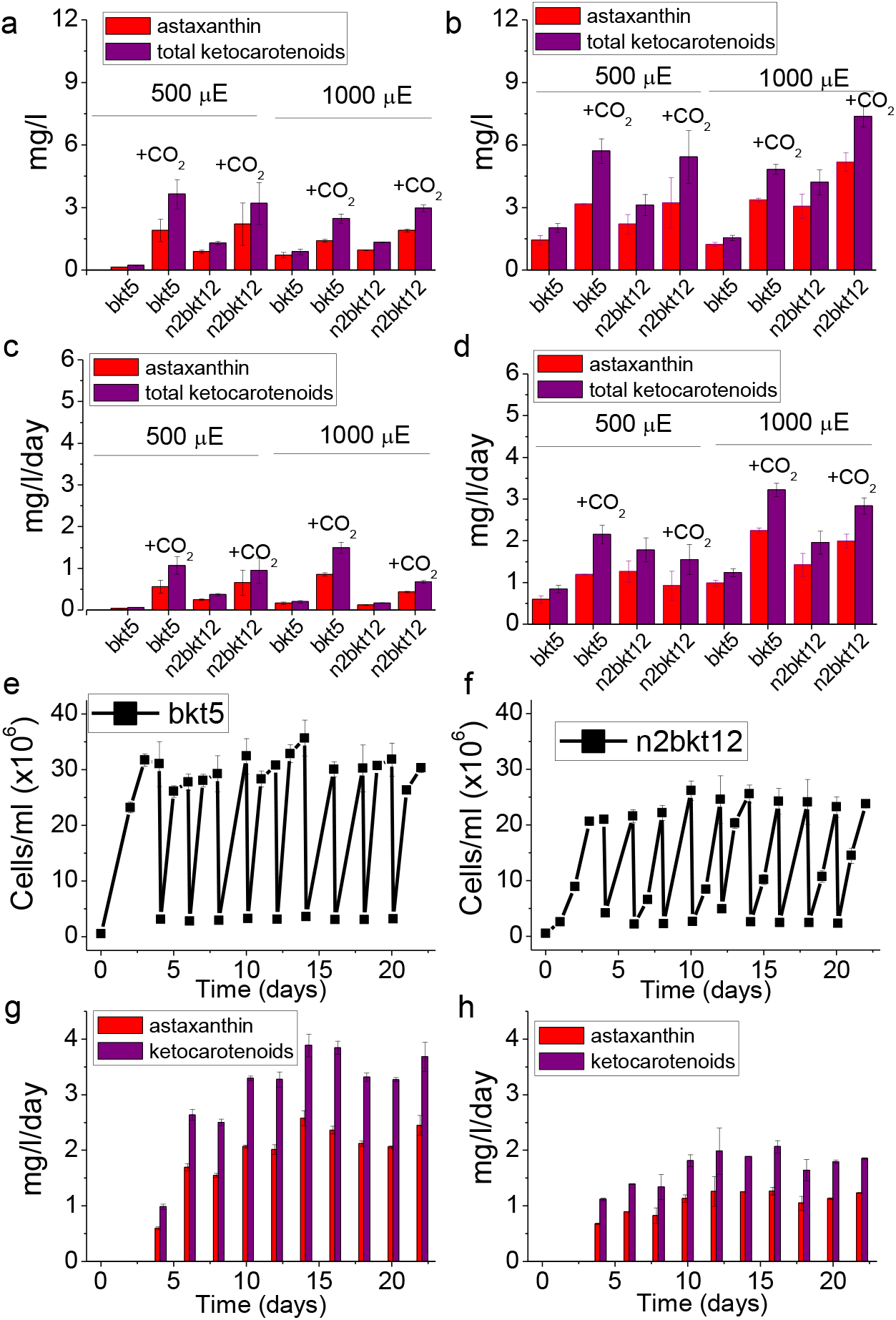
Astaxanthin and ketocarotenoid productivity in different growth conditions. Productivities of astaxanthin (red) and ketocarotenoids (purple) are presented as volumetric (mg L^-1^) **(a, b)** and daily volumetric (mg L^-1^d^-1^) **(c, d)** obtained from *bkt5* and *n2bkt12* mutants in HS (a) or TAP **(b)**. Cells were grown at 500 or 1000 μmol photons m^-2^ s^-1^ with air bubbling or air plus 3% CO_2_ bubbling (CO_2_). Data were obtained by HPLC analysis. **(d, e)** Astaxanthin productivity in semi-continous scale-up experiment: *bkt5* **(d)** or *n2bkt12* **(e)** strains were cultivated in semi-continous mode in 500 ml flasks at 500 μmol photons m^-2^ s^-1^ in TAP with 3% CO_2_ bubbling and stirring. Cells were manually counted and diluted tenfold where they reached stationary phase. **(f)** Astaxanthin (red) and ketocarotenoids (purple) productivities obtained in semi-continuous cultivation, as determined by HPLC analysis.

In general for both strains, mixotrophy was found to result in higher productivities compared to autotrophy (Figure 4a-d). Supplying CO_2_ to the culture clearly increased the productivity in all the different conditions especially when acetate was present (Figure 4a-d). For example, *bkt5* grown in TAP exhibited approximately double productivity rates of astaxanthin and ketocarotenoids when CO_2_ was added to TAP cultures at 500 and 1000 μmol m^-2^ s^-1^ (Figure 4d). Light was observed to be important for the accumulation of astaxanthin in both *bkt5* and *n2bkt12* strains, overall ketocarotenoid production increases were observed at 1000 μmol m^-2^ s^-1^ compared to cells grown at 500 μmol m^-2^ s^-1^ (Figure 4). The highest volumetric productivities obtained in these experiments were obtained for *bkt5* grown in TAP at 1000 μmol m^-2^ s^-1^ with CO_2_ which produced ~2.24 mg ± 0.06 mg astaxanthin L^-1^ d^-1^ and 3.22 ± 0.16 mg total ketocarotenoids L^-1^ d^-1^. The productivities in mg L^-1^ d^-1^ observed for *n2bkt12* were similar compared to the *bkt5* in these conditions (Figure 4d).

An additional experiment was performed in 600 ml stirred flasks with TAP at 500 μmol m^-2^ s^-1^, in order to study the scalability of the productivity obtained with this mutant. The experiment was conducted for 3 weeks in continuous light (Figure 4e, f) and again the productivity of astaxanthin and total ketocarotenoids were determined from the system during the cultivations (Figure 4g,h). Pigment productivities were quantified at every dilution, which was performed when the cells reached stationary phase. Both trains exhibited an adaption phase in the first days of growth in this system, followed by a phase where repetitive dilutions could be maintained for 3 weeks (Figure 4e, f). For strain *bkt.* productivity values reached 3.89 ± 0.19 mg ketocarotenoids L^-1^ d^-1^ with an average of 2.5 ± 0.18 mg astaxanthin L^-1^ d^-1^ (Figure 4g). A slower growth rate of *n2bkt12* led to a lower overall yield of ketocarotenoids and astaxanthin (Figure 4h). Although *n2bkt12* exhibited increase astaxanthin and ketocarotenoid productivity per cell, its slower growth rate hindered its competitiveness with the more rapid-growing *bkt* line. Therefore, further productivity analyses focused only on *bkt5.*

The productivity of astaxanthin and ketocarotenoid of the *bkt* strain was further investigated in 80 ml airlift photobioreactors using conditions of autotrophy or mixotrophy, in presence of 3% CO_2_, and with increasing irradiance up to 3000 μmol m^-1^ s^-1^ (Figure 5). In autotrophy, astaxanthin accumulation was reduced when irradiance was increased past 500 μmol m^-1^ s^-1^, total ketocarotenoid amounts increased, suggesting different carotenogenesis dynamics in very high light conditions (Figure 5). In mixotrophy, both astaxanthin and total ketocarotenoid accumulation were higher compared to autotrophy conditions and partitioning between these carotenoid types was largely conserved across all irradiances (Figure 5C). Astaxanthin and ketocarotenoid productivity was similar at the different irradiances under autotrophic conditions: ~1 mg astaxanthin L^-1^ d^-1^ and ~1,6 mg ketocarotenoids L^-1^ d^-1^. Mixotrophy was found to yield ~ 3,2 mg astaxanthin L^-1^ d^-1^ and ~ 4,1 mg ketocarotenoid L^-1^ d^-1^ at irradiances between 1500-3000 μmol m^-1^ s^-1^.

**Figure 5.**
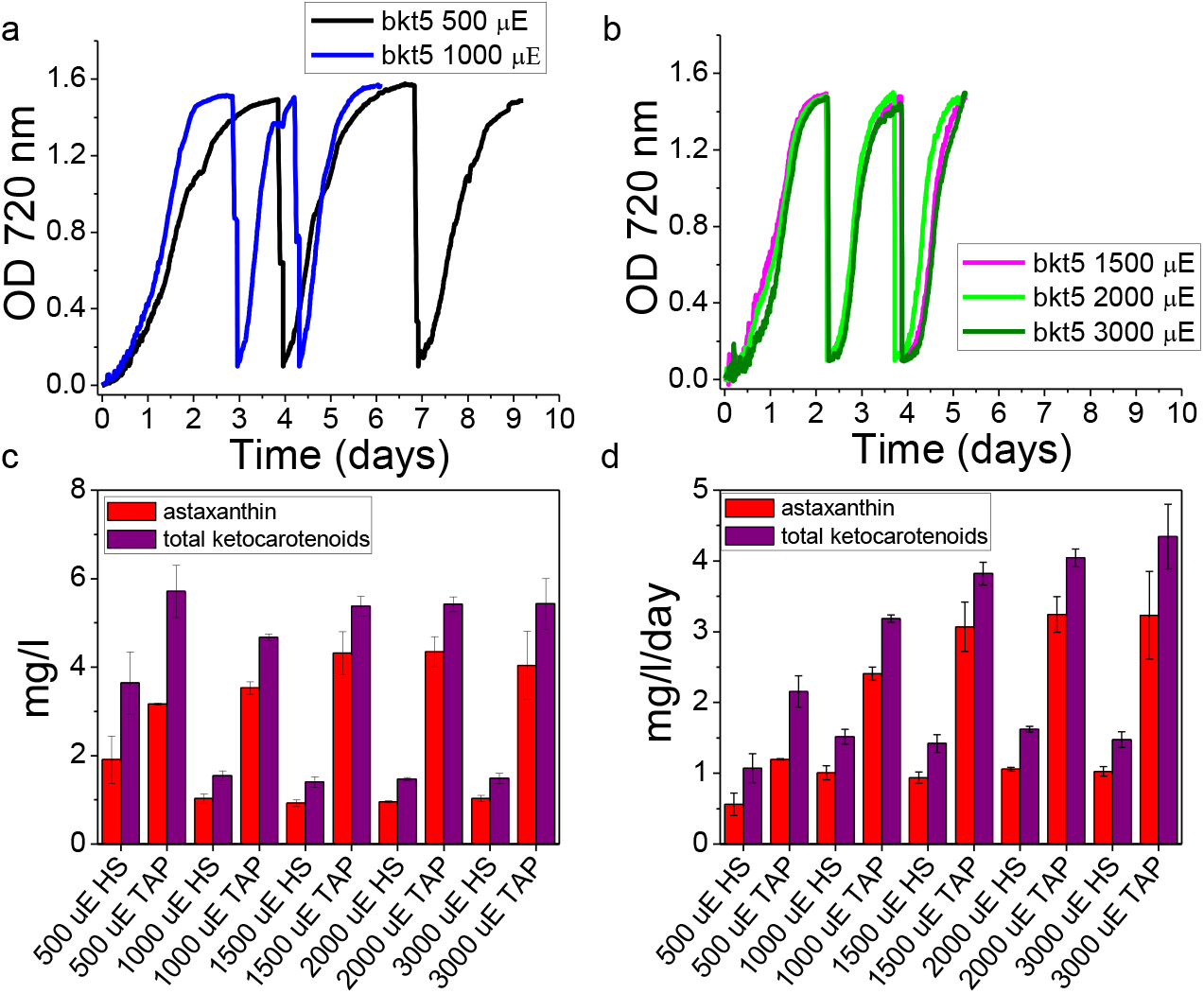
Astaxanthin and ketocarotenoids production by *bkt* lines in very high light conditions. (a, b) Growth curves of bkt lines cultivated with 3% CO_2_ bubbling at 500, 1000 (1000), 1500 (1500), 2000 (2000) or 3000 (3000) μmol photons m^-2^ s^-1^ in mixotrophy conditions (TAP medium). Cells were manually diluted tenfold when the stationary phase was reached. Volumetric **(c)** and volumetric per day **(d)** productivities of astaxanthin (red) and ketocarotenoids (purple) obtained from *bkt5* mutant grown with 3% CO_2_ bubbling at 1000 (1000), 1500 (1500), 2000 (2000) or 3000 (3000) μmol photons m^-2^ s^-1^ in HS or TAP. Data are expressed as means ± SD (n = 4).

### Extractability and bio-accessibility of ketocarotenoids from cells

One of the limitations in natural astaxanthin production from *H. lacustris* microalgae is the difficulty of pigment extraction from the cells which is hindered by the tough cell wall of the aplonospore cysts. These walls also limit the bio-accessibility of *H. lacustris* astaxanthin as they are largely resistant to digestion (Sommer et al., 1991). In order to evaluate a possible benefit in using *C. reinhardtii* for astaxanthin production over *H. lacustris,* the extractability of carotenoids from *C. reinhardtii bkt5* and *H. lacustris* cyst cells was compared by treating cells with solvent (ethyl acetate) or with mineral oil, which is a Generally Recognized as Safe (GRAS) agent (Figure 6). Extraction with dimethyl sulfoxide (DMSO) was also used as a control for these agents as it has been previously reported to be the most effective method for total pigment extraction from *H. lacustris* (Zhekisheva et al., 2002). Treating *H. lacustris* cells with either ethyl acetate or mineral oil gave an extremely low efficiency of astaxanthin extraction; less than 2% of total pigments could be extracted with these agents (Figure 6a,b). With *C. reinhardtii bkt5,* however, ethyl acetate extracted all pigments from *C. reinhardtii* cells and the treatment with mineral oil extracted more than 80% of the carotenoids (Figure 6a, b). These experiments confirmed that extractability of pigments from the transformed *C. reinhardtii* cells is more readily achieved than from *H. lacustris*. Finally, the bio-accessibility of astaxanthin produced in *C. reinhardtii* was compared to *H. lacustris* by simulating gastro-intestinal digestion *in vitro* (Minekus et al., 2014) (Figure 6b, c). Cells were treated with simulated digestion fluids containing buffered solutions and enzymes, i.e. pepsin for the gastric phase and pancreatin for the intestinal phase. (After three hours, almost no pigment was extracted from *H. pluvialis* cysts by this method while more than 80% of the pigments were extracted from *C. reinhardtii bkt5* (Figure 6b). These results indicate that cell wall deficient *C. reinhardtii* is a promising host organism for astaxanthin production as it is readily digestible and, consequently, the produced astaxanthin is more bio-accessible.

**Figure 6.**
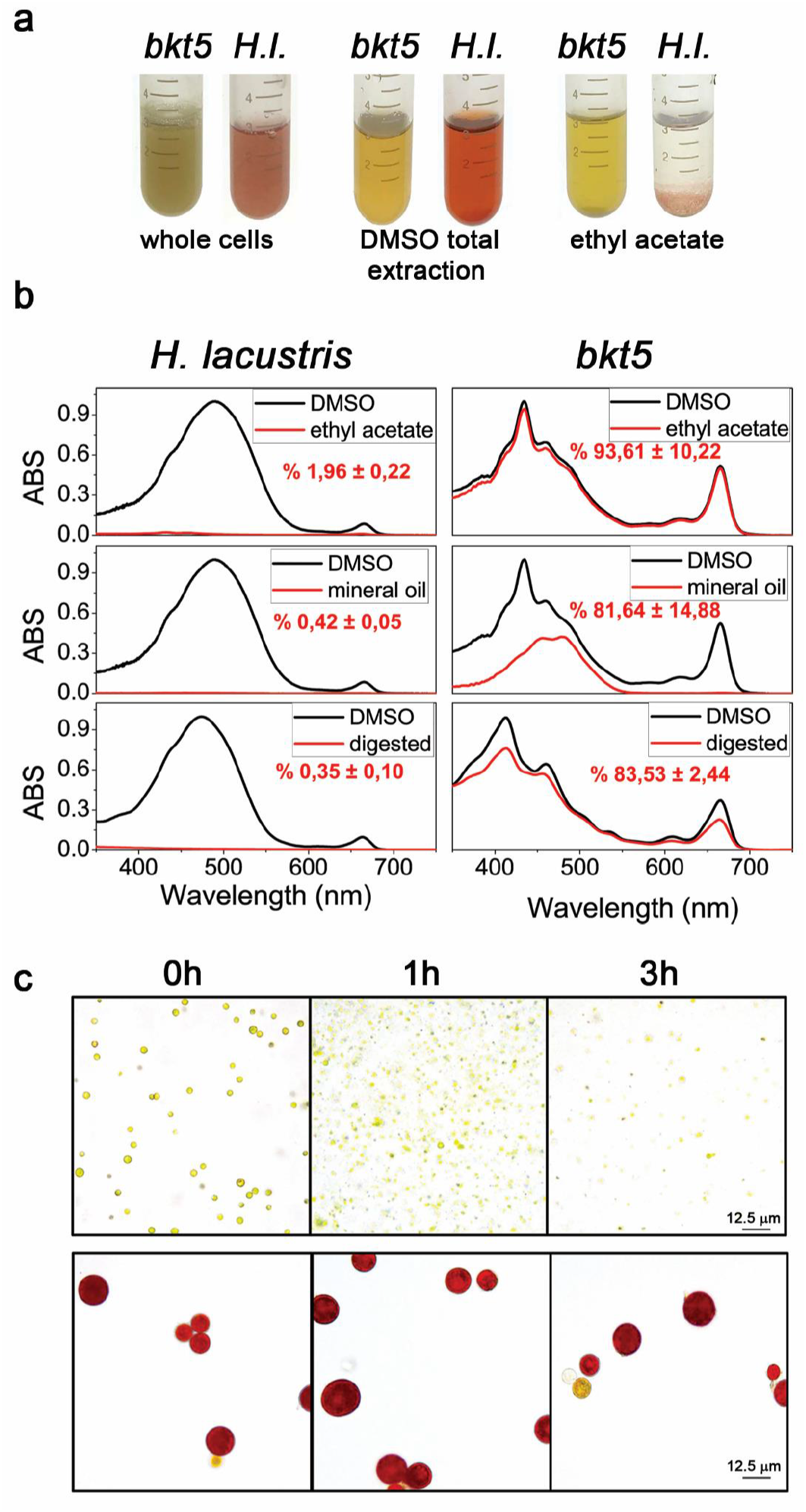
Extractability and bio-accessibility of astaxanthin from *H. lacustris* and *C. reinhardtii* cells. (a) Aplanospore cysts of *H. lacustris* in ‘red phase’ and *C. reinhardtii bkt5* in liquid medium (left) were compared using different pigment extraction methods. DMSO is used as a control from total extraction of pigments from both hosts (center) and ethyl acetate (right) or mineral oil were used. **(b)** Absorption spectra of pigments extracted from *H. lacustris* in red phase and *bkt5* with different solvents. Black traces are the absorption spectra of pigments obtained upon extraction in DMSO (positive control: maximum extraction); red traces show the pigments extracted with different methods. Spectra are normalized to the maximum absorption of the DMSO control. Inserts indicate the percentage of pigments extracted. Data are expressed as mean ± s.d. (n = 3). **(c)** Images of cells of *bkt5* and *H. lacustris* before (0h) and after simulated gastro-intestinal digestion (1 or 3h).

## Discussion

The green microalga *C. reinhardtii* does not naturally accumulate ketocarotenoids, however, a beta-carotene ketolase (CrBKT) is found in its genome. CrBKT has been shown to catalyse the conversion of carotenes into ketocarotenoids when heterologous expression of this gene was performed in other organisms, however, this sequence shows almost no detectible expression in the algal host itself (Figure 1b). Compared to BKT from other organisms, the CrBKT contains a long peptide extension on its C-terminus (Figure 1c), which appears to be an unfolded peptide region. Here, we took the CrBKT sequence and used our recently developed transgene optimization strategy (Baier et al., 2018b) to enable its re-integration into the algal nuclear genome and overexpression. When CrBKT was expressed, it caused the green algal cells to noticeably change colour from green to red, indicating the conversion of carotenoids to ketocarotenoids, with astaxanthin representing a major component of these. We found the best combination of fusion partners and targeting peptides to be a fusion of the native gene, including its chloroplast targeting peptide, with the PsaD chloroplast targeting peptide and replacement of the C-terminal amino acid extension with a GSG-linker and YFP reporter (Figure 2c). The 116 C-terminal amino acid extension of the CrBKT reduced overall astaxanthin productivity when included in expression constructs, however, did not completely abolish astaxanthin formation (Figure 2c). This suggests that the presence of this peptide alone, is not responsible for its lack of expression in the algal host and that its low transcription rates, likely play a larger role. As the presence of astaxanthin did not perturb growth rates of either UVM4-bkt or npq2-bkt expression strains, at least in the conditions tested here (Figure 3), it is unclear what advantage repression of this native gene has given to the algal host. It is possible that a condition could exist where it is expressed, indeed, *Chlamydomonas nivalis,* a relative of *C. reinhardtii,* is known to produce astaxanthin as a photoprotective pigment under variable high-light conditions on snow (Rezanka et al., 2008). However, we did not find a condition where this gene was expressed to notable levels (based on FPKM) in the transcriptome databases investigated here (Figure 1b). Due to the very low rates of transcription, and the clear lack of ketocarotenoids in *C. reinhardtii*, we have called *Cr*BKT a pseudogene in this work.

Astaxanthin biosynthesis requires both a ketolase (BKT) and a hydroxylase (CHYB) (Figure 1a). A *CHYB* gene is present in the nuclear genome of *C. reinhardtii* (Cre04.g215050) and it is expressed, even if to a lower extent compared to other genes related to carotenoid biosynthesis (Figure 1) (Lohr et al., 2005). Re-insertion of only CrBKT, into the nuclear genome of *C. reinhardtii* allowed the synthesis of high amounts of astaxanthin and other ketocarotenoids that normally do not accumulate in this organism (Figure 2). These findings indicate that the native *Cr*CHYB was highly functional and able to participate in ketocarotenoid biosynthesis. When *Cr*BKT was overexpressed in the alga, astaxanthin became the major carotenoid (up to 50% of total carotenoids) with ketocarotenoids corresponding to ~ 70% of total carotenoids (Table 1). As the major substrate of CrBKT is zeaxanthin, CrBKT overexpression was also attempted in the *npq2* mutant, which is deficient in the zeaxanthin epoxidase and accumulates this as a terminal carotenoid (Niyogi et al., 1997). Indeed, astaxanthin accumulation per cell was higher in *npq2* than UVM4 derived BKT overexpression strains (Table 1). However, *npq2* exhibits a slower growth rate than UVM4 (Figure 3), and although the growth was not affected by the presence of astaxanthin, overall volumetric productivities were higher with faster-growing UVM4 derived *bkt* strains (Figure 4).

The *bkt* mutants show a strong reduction of chlorophyll content of ~ 80% with respect to the parental line although remain photosynthetically active (Table 1, Figure 3b, Figure 4). The xanthophylls lutein, violaxanthin and neoxanthin are ligands for the photosystem antenna subunits, β-carotene is a ligand for both photosystem core complexes and antenna, while astaxanthin, in *H. lacustris*, is manly accumulated as a free form in the membrane, although it has been reported to bind photosystems (Mascia et al., 2017). In a photosynthetic cell, the absence of carotenoids hinders photosystem assembly and reduces chlorophyll content (Cazzaniga et al., 2016). It is possible that by rapidly converting carotenoids into ketocarotenoids, photosystem assembly is hindered, and lower chlorophyll amounts accumulate in the algal cell. Another possible reason for the observed decrease of chlorophyll contents in cells expressing CrBKT could be that photosynthetic complexes, assembled without proper carotenoids, generate more ROS that damage photosynthetic membrane (Cazzaniga et al., 2016), however, this hypothesis is less probable as not perturbation in growth rates were observed. In addition, high-light intensities were tolerated by cells, and ketocarotenoids exhibit high antioxidant capacities which would likely protect membranes from ROS damage. In engineered plants, strains with an almost complete conversion of carotenoid to ketocarotenoids have been generated and showed slower growth rates, reduced photosynthetic parameters and increased photoinhibiton (Fujii et al., 2016; Hasunuma et al., 2008; Zhong et al., 2011). In *C. reinhardtii bkt* expression strains, the 30% of standard carotenoid and 20% of the standard chlorophyll were left in engineered strains and seemingly were enough to enable light energy absorption needed photosynthetic growth. The reddish-brown phenotype (Figure 3a) of *bkt* transformants could be interesting for industrial application, as they are characterized by a decreased chlorophyll content which may allow better light penetration into dense algal cultures. Indeed, reduced antennae and chlorophyll containing phenotypes have previously been reported to promote increased algal culture productivity (Cazzaniga et al., 2014; Melis, 2009).

*H. pluvialis* is currently considered the best natural source and the main commercial producing organism for astaxanthin as this alga can accumulate up to 4% of its dry weight as astaxanthin (Boussiba and Vonshak, 1991). Although *C. reinhardtii* strains presented in this work accumulate ketocarotenoids up to only 0.2 % of their biomass, by investigating even minimal optimization of growth conditions, productivities of ketocarotenoids could be reached up to ~ 4,5 mg L^-1^ d^-1^. Due to the need of a two-phase growth and astaxanthin induction, as well as a strong recalcitrant cell wall of astaxanthin containing aplanospores, the yield of astaxanthin from *H. lacustris* has been reported from 0.12 to 15 mg L^-1^ d^-1^ (Lopez et al., 2006; Park et al., 2014). Therefore, ketocarotenoid production for *C. reinhardtii bkt* mutants is within in the same range of that of *H. lacustris*, however, requiring significantly few process parameters or extraction techniques. Indeed, further optimization of *C. reinhardtii* cultivation, for example, in higher-density reactor concepts, will likely improve this efficiency.

Different techniques have been developed to disrupt *H. lacustris* cell walls and recover astaxanthin, these involve mechanical processes like high pressure or bead milling as well as the use of solvents, supercritical carbon dioxide, or enzymatic digestion (Shah et al., 2016). All these procedures increase the cost of astaxanthin bio-production processes and exhibit a variable range of extraction efficiencies. *C. reinhardtii* strains commonly used for transformation and expression experiments generally lack cell walls. These strains still produce cell wall proteins, however, they are not able to assemble them on the cell surface and the proteinaceous wall proteins are instead secreted into the culture medium (Baier et al., 2018a). Here, we used the UVM4 strain which is a derivative of the *cw15* line of cell-wall deficient mutants (Neupert et al., 2009). Due to the reduced cell wall, this strain is readily disrupted, enhancing the relative ease of pigment extraction from *bkt* expression lines. Astaxanthin extractability of *H. lacustris* aplanospores using ethyl acetate or mineral oil was below 1% while pigments in *C. reinhardtii bkt* lines could be completely extracted under similar conditions (Figure 6). This increased extractability also means that pigments within engineered *C. reinhardtii* are more bio-accessible, as the cyst state of astaxanthin containing *H. lacustris* may not be readily digested in target organisms (fish, livestock, or humans). *In vitro* simulated digestion showed enhanced pigment extraction from C. reinhardtii bkt5 compared to *H. lacustris*, indicating the engineered alga could be used directly in aquaculture and nutraceutical feed, without the need for prior pigment extraction (Figure 6b). Direct used of algal biomass as a source of ketocarotenoid pigments for livestock and aquaculture could also benefit the quality of pigments delivered to these organisms as storage of carotenoids can lead to their degradation or oxidation.

Given it only requires the overexpression of a single ketolase to generate constitutive astaxanthin and ketocarotenoid generation in *C. reinhardtii*, it will likely be possible to transfer this biosynthesis rapidly into other more industrially cultivated algal strains. For example, certain *Chlorella* species have demonstrated robust outdoor growth, or the genetically amenable *Nannochloropsis* sp. may be alternatives where the production of astaxanthin can be scaled in existing infrastructures (Liu et al., 2014; Lubián et al., 2000). However, both of these algae also exhibit robust cell walls, which may limit overall process productivity. Indeed, *C. reinhardtii* has been shown amenable to scale up in air-lift bioreactors even under outdoor conditions (Lauersen, 2018) and it may be that cultivation of cell-wall deficient astaxanthin producing *C. reinhardtii* presents the best case of rapid growth and ease of pigment extractability. Given the robust activity of the *Cr*BKT, it may be likely that further customizations of the carotenoid biosynthesis can be readily achieved in this alga. Future engineering targets may seek to use existing carotenoids, or these new populations of ketocarotenoids, as even further targets for bioengineering optimization in this versatile algal host.

## Experimental procedures

### Algal cultivation and strain maintenance

*C. reinhardtii strain* UVM4 *was* graciously provided by Prof. Dr. Ralph Bock and *npq2* (CC-4101) was obtained by Chlamydomonas Resource Center (https://www.chlamycollection.org). Both strains were maintained on TAP agar plates or in liquid shake flasks at 25 °C with 100-150 μmol photons m^-2^s^-1^ of continuous white light. Growth tests were conducted using different systems: shaking flasks, stirring flasks or Multi-Cultivator MC-1000 (Photon Systems Instruments). Temperature was controlled to 25 °C while light intensities were varied as indicated in the text. Tris-Acetate-Phosphate (TAP) or High Salts minimal (HS) media were used for mixotrophic or photoautotrophic conditions, respectively as described in the text (Harris and Harris, 2008).

### Design and cloning of expression cassettes

The BKT (AY860820.1) found within the genome of *C. reinhardtii* was synthetically redesigned with codon optimization and intron spreading as recently described (Baier et al., 2018b). The synthetic optimized CrBKT gene coding sequence (CDS) was deprived of the last 345 bp and a small region was added to make a C-terminal GSG linker prior to optimization and the synthetic gene produced by GeneArt (Germany). The synthetic CrBKT sequence was cloned between *Bam*HI-*BglII* into the pOpt2_PsaD_mVenus_Paro vector (Wichmann et al., 2018) to generate a protein which contains the *C. reinhardtii* photosystem I reaction centre subunit II (PsaD) chloroplast targeting peptide and a C-terminal mVenus (YFP) fusion. The emptry vector served as a control. The vector confers resistance to paromomycin from a second expression cassette (Lauersen et al., 2015). Variations on this construct were generated by successive cloning within the pOpt2_PsaD_CrBKT_YFP_Paro vector. The CrBKT was cloned into the same vector but without PsaD target peptide in order to obtain a BKT protein targeted with endogenous transit peptide. To test the C-terminal amino acid extension, the whole BKT protein was generated by PCR fusion of a 345 bp region to its C-terminus. First, the C-terminal region was chemically synthesized including last 30bp of BKT previously generated in order to allow primer binding. Then two different fragments were amplified: one containing BKT sequence using pOpt2_PsaD_BKT_YFP as template (primer: for: 5’-GGCCGGATCCGGCCCCGGCATCCAGCCCACCAGCG-3’, rev: 5’-CCGCGGTCGCGGCAGCCGCGGCGGGCACGGGCAGGGGGCCGGGGGCCAGGGCGGCGCCGCGG GCGATC-3’) while the second contained the BKT C-terminal region using synthetic sequence as template (primer for: 5’-GATCGCCCGCGGCGCCGCCCTG-3’, rev: 5’-GGCCAGATCTGCCGCTGCCGGCCATCACGCCCACGGGGGCCAGC-3’). The two fragments were then used as template for an additional amplification to fuse these elements using additional primers for: 5’-GGCCGGATCCGGCCCCGGCATCCAGCCCACCAGCG-3’ and rev: 5’-GGCCAGATCTGCCGCTGCCGGCCATCACGCCCACGGGGGCCAGC-3’. The amplified sequence was cloned into pOpt2_PsaD_BKT_YFP_Paro in the *Bam*HI-*Bgl*II position.

pOpt2_PsaD_BKT without YFP was generated by removing YFP sequence using ZraI and EcoRV enzymes from pOpt2_PsaD_BKT_YFP. pOpt_PsaD_BKT2x_YFP was generated by first PCR amplification of BKT sequence to remove the 5’ 150 nucleotides coding for the N-terminal targeting peptide using pOpt2_PsaD_BKT_YFP_Paro as template (primer: for 5’-GGCCGGATCCAAGCTGTGGCAGCGCCAGTACCACCTG-3’, rev: 5’-GGCCAGATCTGCCGCTGCCGGCCATCACGCCCACGGGGGCCAGC-3’). This sequence was inserted into a pOpt2 vector generating pOpt2_PsaD_-50aaBKT_YFP which could then be combined with the pOpt2_PsaD_BKT_YFP_Paro vector by compatible BamHI-BglII overhangs to generate the pOpt2_PsaD_2xBKT_YFP_Paro. For subcellular localization determination mediated of the BKT N-terminal chloroplast targeting peptide, 102 and 120 bp from the 5’ region of the BKT coding for its N-terminus was amplified and cloned into *NdeI-BglII* sites of the pOpt2_PsaD_BKT_YFP_Paro (primer for: 5’-GGCCGGATCCGGCCCCGGCATCCAGCCCACCAGCG-3’ rev:amino acid 34 for 102bp and rev:amino acid 40 for 120bp).

All cloning described here was performed using FastDigest restriction enzymes (Thermo Scientific) and the Rapid DNA Dephos & Ligation Kit (Roche) following manufacturer’s instructions. PCR was conducted using Q5^®^ High Fidelity DNAPolymerase (NEB). DNA fragments were separated in 2% (w/v) agarose gels and purified using the peqGOLD Gel Extraction Kit (VWR). Heat-shock transformation of chemically competent *Escherichia coli* DH5α cells were performed for all vectors followed by selection on LB-agar plates with ampicillin as a selection marker. Colonies were evaluated by colony PCR followed by plasmid isolation from overnight culture using peqGOLD Plasmid Miniprep Kit I (VWR). All sequences were confirmed by Sanger sequencing (Sequencing Core Facility, CeBiTec, Bielefeld University).

### C. reinhardtii transformation and mutant screening

Nuclear transformation was carried out by glass beads agitation as previously described (Kindle, 1990) using 10μg of linearized plasmid DNA. Selection of transformants was done on TAP agar plates supplied by paromomycin (10 mg l^-1^) for 5-7 days. Positive expressing colonies were pre-screened visually for red colouration before further quantification.

### Fluorescence microscopy localization

YFP fluorescence imaging was performed as previously described (Lauersen et al., 2016).

### Growth analysis

Parameters used for monitor growth were cell density and cell dry mass. Cell densities were measured using Countess II FL Automated Cell Counter (Thermo Fisher Scientific). Dry biomass was evaluated by overnight lyophilization of washed cell pellets and gravimetric determination.

### Pigment analyses

Pigments were extracted from intact cells using 80% acetone buffered with Na2CO3 and analyzed by absorption spectra followed by curve fitting or by reverse phase HPLC. Absorption spectra were measured with Jasco V-550 UV/VIS spectrophotometer as described in (Cinque et al., 2000). Spectra were fitted as described in (Croce et al., 2002) introducing in the fitting method the astaxanthin absorption form in acetone 80%: considering the similar absorption of astaxanthin and cantaxanthin, the results obtained by fitting of pigment extracts absorption spectra were considered as representative to total ketocarotenoids. Reverse phase HPLC was conducted as described in (Scibilia et al., 2015). Briefly, Thermo-Fisher HPLC system equipped with a C18 column using a 15-min gradient of ethyl acetate (0 to 100%) in acetonitrile-water-triethylamine (9:1:0.01, vol/vol/vol) at a flow rate of 1.5 ml/min was used. Pigment detection was conducted with a Thermo-Fisher 350−750nm diode array detector. Astaxanthin and cantaxanthin peaks were identified by comparison to commercially available standards (CaroteNature GmbHas).

### Extractability and simulated digestion of ketocarotenoids in H. lacustris and C: reinhardtii

Same weight of dried cells of *H. lacustris* and *bkt5* were resuspended in water and treated with ethyl acetate or mineral oil for 20 min at room temperature and subsequently subject to centrifugation. Extracted pigments present in the supernatant were recovered and spectra were recorded to determine relative extractability. Treatment was repeated once for ethyl acetate and three times for mineral oil. The simulate digestion was performed following the protocol described by (Minekus et al., 2014) with some modification. 0,1 gr of freeze-dried cells (LIO-5P, 5pascal) were resuspended in 1 ml of simulated gastric fluid and stirred for 1 hour at 37° C, then 2 ml of simulated intestinal fluid was added to the samples. After further 2 hours at as above, samples were centrifuged for 3 minutes at 3000 g to pellet intact cells and isolate the supernatant digested fraction. Pigments were extracted with acetone from the digested fraction and spectra recorded.

## Acknowledgements

The authors would like to acknowledge support of the technology platform and infrastructure at the Center for Biotechnology (CeBiTec) of Bielefeld University (T.B., K.J.L., L.W.). The research was supported by the ERC Starting Grant SOLENALGAE (679814) to M.B. Authors declare no conflict of interest.

## Short legends for Supporting Information

Figure S1. Western blot and immunodetection of BKT-YFP fusions

Figure S2. Microscopy images of *H. lacustris* and *C. reinhardtii* BKT overexpressing strain (*bkt* 5).

