## Supplementary information for "Turning a green alga red: engineering astaxanthin biosynthesis by intragenic pseudogene revival in *Chlamydomonas reinhardtii*"

**Figure S1. Western blot and immunodetection of BKT-YFP fusions.** Proteins were separated by 12% Tris-glycine-SDS-PAGE. Separated proteins were stained using colloidal Coomassie. Brilliant Blue G-250 or analyzed by immunodetection on nitrocellulose membranes using a HRP-linked rabbit-anti-GFP antibody (Thermo Scientific, A10260) and Pierce™ ECL Western Blotting substrates (Thermo Scientific).

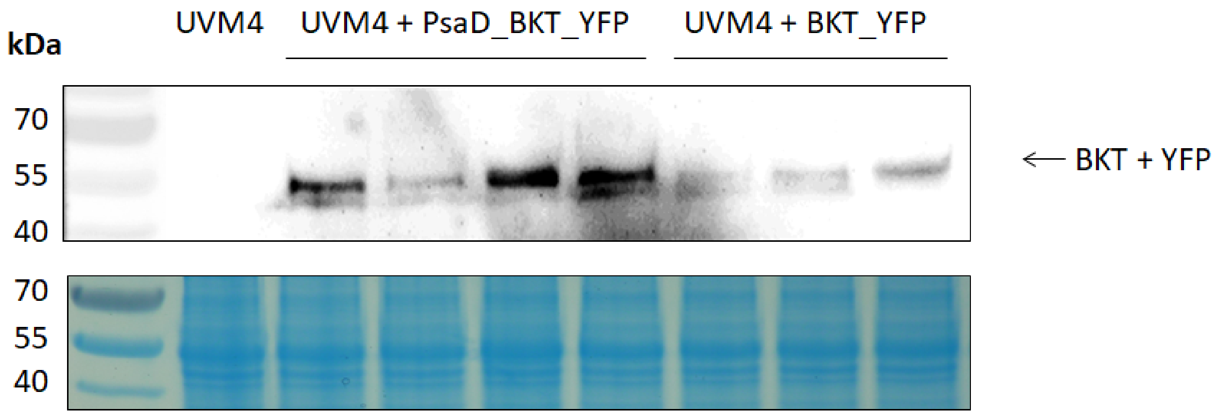

**Figure S2. Microscopy images of *H. lacustris* and *C. reinhardtii* BKT overexpressing strain (*bkt* 5).** *H. lacustris* cells are presented in green and red phases. Red phase was obtained by high light treatment and nitrogen starvation for 5 days.

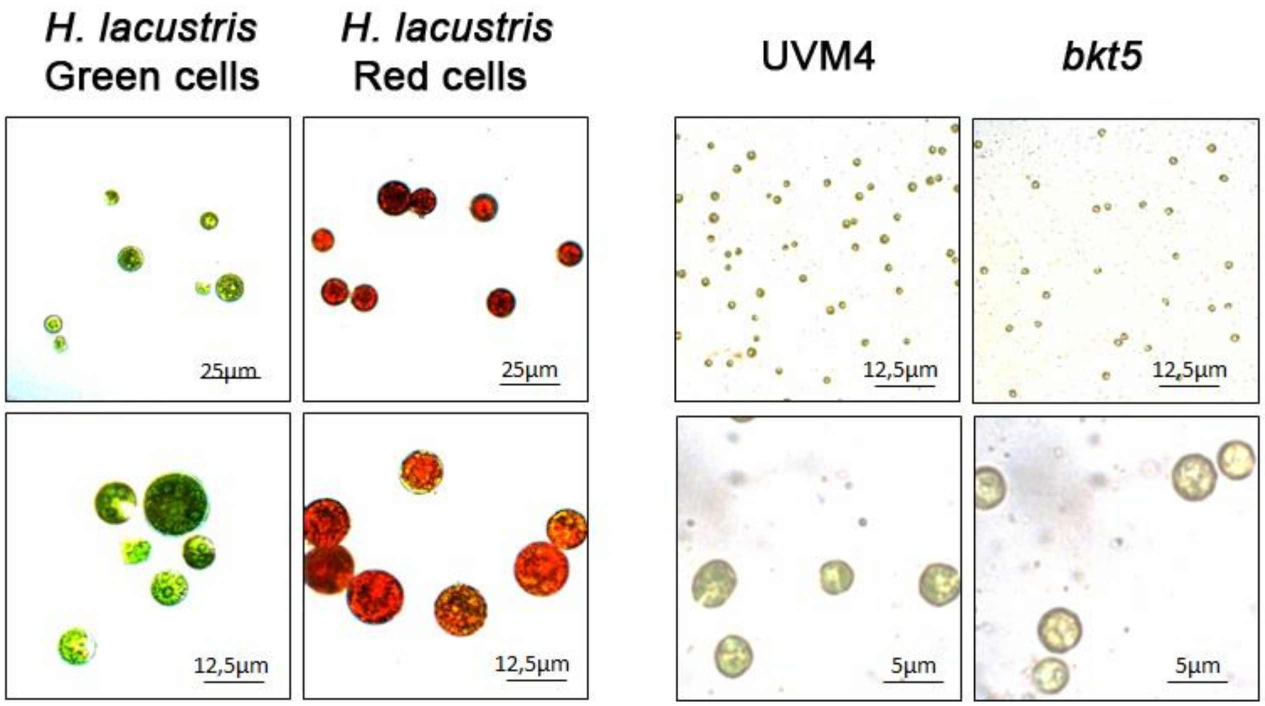
